# Semaphorin4D induces inhibitory synapse formation by rapid stabilization of presynaptic boutons via MET co-activation

**DOI:** 10.1101/100271

**Authors:** Cátia P. Frias, Tom Bresser, Lisa Scheefhals, Hai Yin Hu, Paul M. P. van Bergen en Henegouwen, Casper C. Hoogenraad, Corette J. Wierenga

**Affiliations:** Cell Biology, Department of Biology, Faculty of Science, Utrecht University, 3584 CH Utrecht, the Netherlands

## Abstract

Changes in inhibitory connections are essential for experience-dependent circuit adaptations. Defects in inhibitory synapses are linked to neurodevelopmental disorders, but the molecular processes underlying inhibitory synapse formation are not well understood. Here we use high resolution two-photon microscopy in organotypic hippocampal slices to examine the signaling pathways induced by the postsynaptic signaling molecule Semaphorin4D (Sema4D) during inhibitory synapse formation. By monitoring changes in individual GFP-labeled presynaptic boutons we found that the primary action of Sema4D is to induce stabilization of presynaptic boutons within tens of minutes. Stabilizing boutons rapidly recruited synaptic vesicles, which was followed by accumulation of postsynaptic gephyrin. Newly formed inhibitory synapses were complete and functional after 24 hours, as determined by electrophysiology and immunohistochemistry. We further showed that Sema4D signaling is regulated by network activity and can induce a local increase in bouton density, suggesting a possible role in circuit adaptation. We further examined the intracellular signaling cascade triggered by Sema4D and found that bouton stabilization occurred through rapid remodeling of actin, and this could be mimicked by the actin-depolymerizing drug Latrunculin B or by reducing ROCK activity. The intracellular signaling cascade required activation of the receptor tyrosine kinase MET, which is a well-known autism risk factor. Our immunohistochemistry data suggests that MET may be localized to presynaptic inhibitory axons. Together, our data yield important insights in the molecular pathway underlying activity-dependent Sema4D-induced synapse formation and reveal a novel role for MET in inhibitory synapses.

**Significance Statement:** GABAergic synapses provide the main inhibitory control of neuronal activity in the brain. We make important steps in unraveling the molecular processes that take place when formation of inhibitory synapses is triggered by a specific signaling molecule, Sema4D. We find that this process depends on network activity and involves specific remodeling of the intracellular actin cytoskeleton. We also reveal a previously unknown role for MET in inhibitory synapses. As defects in GABAergic synapses have been implied in many brain disorders, and mutations in MET are strong risk factors for autism, our findings urge for a further investigation of the role of MET at inhibitory synapses.

## INTRODUCTION

GABAergic synapses provide the main inhibitory control over neuronal activity in the brain and are indispensable for shaping network function (Isaacson and Scanziani, 2011). In postnatal brain tissue, in which the majority of inhibitory connections have been established, synapse formation and disassembly is still ongoing (Caroni et al., 2012). Formation and disassembly of inhibitory synapses in the brain play an important role in experience-dependent circuit adaptation (Hensch, 2005; Keck et al., 2011; Chen et al., 2015; Froemke, 2015; Sprekeler, 2017) and defects in GABAergic synapses have been observed in many neurodevelopmental disorders (Marín, 2012; Cellot and Cherubini, 2014; Nelson and Valakh, 2015). We and others have previously shown that inhibitory axons are dynamic structures with boutons forming and disappearing with apparently stochastic dynamics (Fu and Huang, 2010; Dobie and Craig, 2011; Kuriu et al., 2012; Frias and Wierenga, 2013). These ongoing bouton dynamics allow quick updating of connections in response to changes in the neuronal circuitry (Staras, 2007; Frias and Wierenga, 2013). New inhibitory synapses form by the emergence of new presynaptic boutons at pre-existing axon-dendrite crossings (Wierenga et al., 2008; Schuemann et al., 2013). However, the signaling pathways that regulate the multiple steps during inhibitory synapse formation are not well understood (Wierenga, 2017).

In the recent years, enormous progress has been made by the identification and characterization of proteins that are involved in the formation of inhibitory synapses (Siddiqui and Craig, 2011; Krueger-Burg et al., 2017; Lu et al., 2017). The class 4 semaphorin Sema4D, originally identified as an axon guidance factor (Kolodkin et al., 1993; Pasterkamp, 2012), has been shown to play a crucial role in this process. Formation of GABAergic synapses was shown to depend on Sema4D signaling, as knockdown of postsynaptic Sema4D led to a 30% reduction of GABAergic synapses in primary cultures (Paradis et al., 2007). In addition, acute activation of Sema4D pathway by adding a soluble form of the extracellular part of Sema4D to primary hippocampal cultures induces rapid increase of GABAergic synapses (Kuzirian et al., 2013; Raissi et al., 2013). The observation that somatic and dendritic synapses responded equally to Sema4D signaling (Kuzirian et al., 2013) suggests that Sema4D could be acting on a broad range of (or perhaps all) inhibitory synapses.

Despite its well-characterized physiological importance, relatively little is known about the cellular mechanism by which Sema4D induces inhibitory synapse formation. It was shown that Sema4D acts as a postsynaptic protein and requires its receptor PlexinB1 to induce inhibitory synapses (Kuzirian et al., 2013; Raissi et al., 2013). The signal cascades that are triggered by Sema4D-PlexinB1 interaction have been well studied in other cells and these studies revealed that Sema4D action is highly cell-specific (Zhou et al., 2008; Cagnoni and Tamagnone, 2014). For instance, Sema4D has been reported to either suppress or enhance cellular adhesion and/or migration, depending on the cell type (Oinuma et al., 2006; Basile et al., 2007; Giacobini et al., 2008; Swiercz et al., 2008). The intracellular molecular events downstream of Sema4D/PlexinB1 signaling that lead to inhibitory synapse induction are currently not known.

In the current study, we used high resolution two-photon microscopy in organotypic hippocampal slices to characterize Sema4D regulation of inhibitory synapse formation in intact tissue and to examine the underlying molecular pathway. We found that Sema4D signaling specifically regulates the rapid stabilization of inhibitory boutons along the axon and that local bouton stabilization by Sema4D can result in local changes in bouton density within tens of minutes. These rapid presynaptic changes are followed by subsequent recruitment of pre-and postsynaptic proteins to complete the formation of functional inhibitory synapses over the course of the next hours. We also found that Sema4D-induced bouton stabilization is activity-dependent. The intracellular pathway for bouton stabilization involves specific remodeling of the actin cytoskeleton, and requires the activation of the receptor tyrosine kinase MET. Our data unravel an important regulatory pathway of activity-dependent inhibitory synapse formation and reveal a novel role for the receptor tyrosine kinase MET in Sema4D-induced formation of inhibitory synapses.

## EXPERIMENTAL PROCEDURES

### Animals

All animal experiments were performed in compliance with the guidelines for the welfare of experimental animals issued by the Federal Government of The Netherlands. All animal experiments were approved by the Animal Ethical Review Committee (DEC) of Utrecht University.

### Hippocampal slice cultures

Hippocampal slice cultures (400 μm thick) were prepared from postnatal day 5-7 of both male and female GAD65-GFP mice (López-Bendito et al., 2004) as previously described (Müllner et al., 2015). In short, the hippocampi were dissected in ice-cold HEPES-GBSS (containing 1.5 mM CaCl_2_·2H_2_O, 0.2 mM KH_2_PO_4_, 0.3 mM MgSO_4_·H_2_O, 5 mM KCl, 1 mM MgCl_2_6H_2_O, 137 mM NaCl, 0.85 mM Na_2_HPO_4_ and 12.5 mM HEPES) supplemented with 1 mM kynurenic acid and 25 mM glucose, and plated in a MEM-based medium (MEM supplemented with 25 % HBSS, 25 % horse serum, 30 mM glucose and 12.5 mM HEPES). In GAD65-GFP mice, approximately 20% of the CA1 interneurons express GFP from early embryonic developmental stage into adulthood (López-Bendito et al., 2004; Wierenga et al., 2010). The majority of GFP-labeled interneurons expresses reelin and VIP, while parvalbumin and somatostatin expression is nearly absent (Wierenga et al., 2010). For our study, the relatively low number of GFP-positive axons is crucial for proper analysis of individual boutons.

The slices were kept in culture for at least one week before the experiments (range 7-21 days *in vitro*) at 35 °C in 5 % CO_2_. For live imaging experiments, slices were transferred to an imaging chamber, where they were continuously perfused with carbogenated artificial cerebrospinal fluid (ACSF; containing 126 mM NaCl, 3 mM KCl, 2.5 mM CaCl_2_, 1.3 mM MgCl_2_, 1.25 mM NaH_2_PO_4_, 26 mM NaHCO_3_, 20 mM glucose and 1mM Trolox). The temperature of the chamber was maintained at 37°C. Treatment and control experiments were conducted in slices from sister cultures.

### Pharmacological treatments

The following drugs were used: 0.1/0.2 % DMSO, 1 nM Fc and Sema4D-Fc (amino acids 24-711) (both R&D Systems), 100 nM Latrunculin B (Santa Cruz Biotechnology), 200 nM Jasplakinolide (Tocris Bioscience), 1 μM PHA-665752 (Sigma-Aldrich) and 10 μM Y-27632 (Sigma-Aldrich). We used the small molecule PHA-665752 (PHA), a highly specific MET inhibitor (Christensen et al., 2003; Deguchi et al., 2016), to decrease endogenous phosphorylation of MET, without affecting MET expression or neuronal cell viability. We used 10 nM Fc or Sema4D for the local puffing experiments.

For treatments that were followed by immunostaining of inhibitory synapses, 1 nM Fc or Sema4D-Fc was added to the culturing medium and slices were left in the incubator for 2, 6 or 24 h before fixation.

### Two-photon imaging

For acute treatments, drugs were added to the perfusion ACSF after a baseline period of 40 minutes (5 time points) and we continued imaging for an additional 10 time points in the wash-in period (total imaging period is 140 minutes). In longer treatments, we treated the slices for 6 hours after the baseline period (5 imaging time points) at the microscope and restarted imaging for 5 time points, for a total treatment period of 6 hours and 40 minutes (400 minutes). For activity blockade, 0.5 μM tetrodotoxin citrate (TTX; Tocris Bioscience) was added to the perfusion ACSF prior to the transfer of the slice to the imaging chamber. Time-lapse two-photon microscopy images were acquired on a Femtonics 2D two-photon laser-scanning microscope (Budapest, Hungary), with a Nikon CFI Apochromat 60X NIR water-immersion objective. GFP was excited using a laser beam tuned to 910 nm (Mai Tai HP, Spectra Physics). The 3D images (93.5 μm x 93.5 μm in xy, 1124 x 1124 pixels) consisted of 29-33 z stacks (0.5 μm step size in z). Small misalignments due to drift were manually compensated during the acquisition.

For the local treatment, we used HEPES-ACSF (containing_126 mM NaCl, 3 mM KCl, 2.5 mM CaCl_2_, 1.3 mM MgCl_2_, 1.25 mM NaH_2_PO_4_, 20 mM glucose, and 10 mM HEPES; pH 7.41) with 20 μM Alexa 568 (Invitrogen), in order to visualize the spread of the local puff. Sema4D or Fc was added to the HEPES-ACSF to a final concentration of 10 nM. The solution was loaded into a patch pipette (4-6 MOhm), and was locally applied to a GFP-labeled axon using a Picospritzer II (General Valve). Time-lapse two photon microscopy imaging was performed as described previously, except that a second laser (Spectra Physics) was used at 840 nm to visualize the area of the puff. The 3D images (51.3 μm x 51.3 μm in xy, 620 x 620 pixels) consisted of 18-22 z stacks (0.5 μm step size in z). After a baseline period of 20 minutes (5 TPs), the pipette was put into position before the stimulation. The stimulation consisted of 300 puffs of 20-50 ms at 2 Hz. The pipette was carefully retracted before continuing the time series for 10 additional time points (total imaging period of 70 minutes).

### Two-photon image analysis

The analysis of inhibitory bouton dynamics was performed semi-automatically using ImageJ (US National Institute of Health) and Matlab-based software (Mathworks). The 3D coordinates of individual axons were selected at every time point by using the CellCounter plugin (Kurt De Vos, University of Sheffield, Academic Neurology). For each image, 1-5 stretches of axons (average length 78 μm with standard deviation 18 μm, with average of 31 boutons per axon with standard deviation 11; for local treatment experiments, average length 39 μm with standard deviation 8 μm, with average of 14 boutons per axon with standard deviation of 4) were selected for analysis.

A 3D intensity profile along the selected axons was constructed at each time point, and individual boutons were identified in a two-step process using custom-made Matlab software (Schuemann et al., 2013). In brief, an axon threshold was calculated to differentiate the axon from the background (2 standard deviations above mean intensity); subsequently, a local threshold (0.5 standard deviation above mean axon intensity) identified the boutons along the selected axon. Only boutons with at least 5 pixels above bouton threshold were included. Each image stack was visually examined, and false positives and negatives were corrected manually. Only raw data was analyzed; images were median-filtered for illustration purposes only.

Boutons were classified as persistent when they were present during all time points, and non-persistent when they were absent during one or more time points during the imaging session. The average fraction of persistent and non-persistent boutons was calculated by normalization to the average number of boutons per axon. To bias our analysis towards synaptic events (Schuemann et al., 2013), we restricted our analysis to boutons that appeared for at least 2 time points at the same location during the imaging period. We verified that our main conclusions did not change when this restriction was released. Based on their presence during baseline and treatment periods, we defined five subgroups of non-persistent boutons: new boutons (not present during baseline), lost boutons (not present during wash-in), stabilizing boutons (non-persistent during baseline, persistent during wash-in), destabilizing boutons (persistent during baseline, non-persistent during wash-in), and transient boutons (non-persistent in baseline and wash-in) (Fig. 1). Average fraction of each subgroup of boutons was normalized to the total average number of non-persistent (NP) boutons per axon. The duration of each bouton was defined as the number of time points present divided by the total number of time points per period. Bouton density was calculated as the average number of boutons at all time points divided by the 3D axon length.

**Figure 1.**
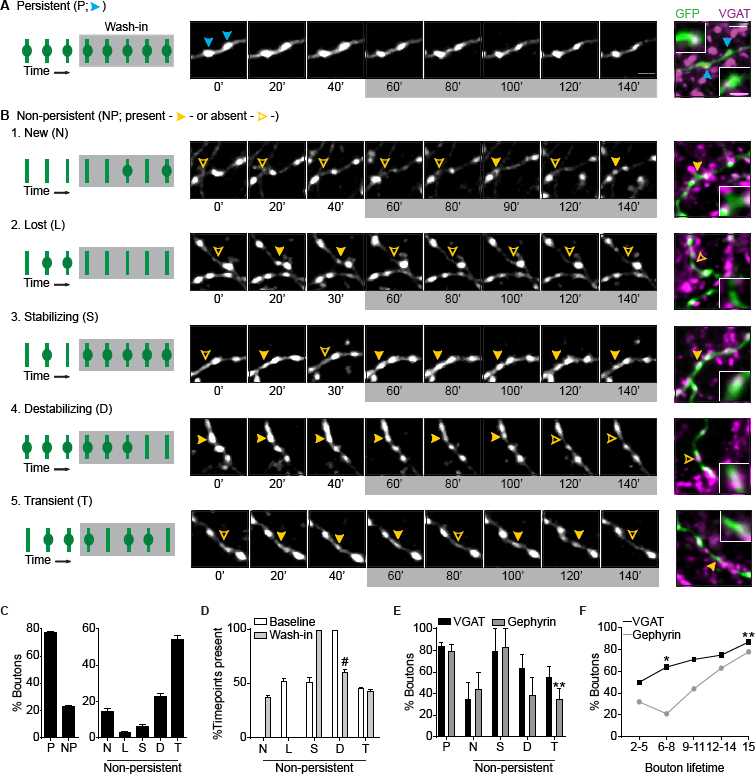
Classification of presynaptic inhibitory boutons by their dynamics. **(A)** Time-lapse two-photon images of two inhibitory boutons (blue arrowheads) along a GAD65-GFP-labeled axon in the CA1 region of the hippocampus. These boutons were present at all time points, and therefore categorized as persistent boutons. Only every second image is shown for clarity. On the right, the same region is shown after fixation and staining against vesicular GABA transporter (VGAT, magenta). The zoom shows a single optical plane through the bouton to demonstrate overlap (white) of VGAT and GFP boutons. Time in minutes. Scale bars 2 μm and 1 μm (zoom).
**(B1-5)** Same as in A, showing examples of new (B1; absent during baseline), lost (B2; absent during wash-in), stabilizing (B3; non-persistent during baseline, and persistent during wash-in), destabilizing (B4; persistent during baseline, and non-persistent during wash-in) and transient (B5; non-persistent during both baseline and wash-in) boutons. Filled yellow arrowheads indicate that the bouton is present, and empty yellow arrowheads indicate that the bouton is absent at the time point shown.
**(C)** Average fraction of persistent (P) and non-persistent (NP) boutons at any given time point, and average fraction of the 5 subgroups of non-persistent boutons normalized to the total number of non-persistent boutons (N - new; L - lost; S - stabilizing; D - destabilizing; T - transient).
**(D)** Percentage of time points in which boutons were present during baseline (white) and wash-in (gray) periods. #: value for D was significantly different from N and T for wash-in (χ^2^; D vs N, *p* = 0.002; D vs T, *p* = 0.002).
**(E)** Fraction of boutons positive for VGAT and gephyrin per axon. Two-way ANOVA analysis showed a significant effect on bouton type (*p* = 0.0098). For gephyrin, P vs T, *p* = 0.001 (Sidak’s multiple comparisons test).
**(F)** Fraction of boutons co-localizing with VGAT or gephyrin as a function of bouton lifetime (total number of time points (#TPs) present during the imaging period). Lost boutons (‘L’ in C,D) were not included. χ^2^: TP2-5, *p* = 0.21; TP6-8, *p* = 0.02; TP9-11, *p* = 0.13; TP12-14, *p* = 0.35: TP15, *p* = 0.008. Confocal images in A are maximum intensity projections of 5-8 z stacks, while two-photon images are maximum intensity projections of 13-15 z stacks. Data are represented as mean ± SEM. Data in C and D from 90 axons from 24 independent experiments, data in E and F from 21 axons from 5 independent experiments for the VGAT staining (P: n=282 boutons; N: n=13; S: n=6; D: n=17; T: n=45) and from 15 axons from 4 independent experiments for the gephyrin staining (P: n=232 boutons; N: n=15; S: n=6; D: n=15; T: n=39). In F, n=14-28 per TP, except for TP15.

### Electrophysiology

During the experiment, the slice was placed in a recording chamber perfused with oxygenated artificial cerebral spinal fluid (ACSF) at a rate of 1 ml/min. The recording ACSF consisted of 126 mM NaCl, 3 mM KCl, 2.5 mM CaCl_2_, 1.3 mM MgCl_2_, 1.25 mM Na_2_H_2_PO_4_, 26 mM NaHCO_3_, and 20 mM glucose. Whole cell voltage clamp recordings were performed at 35 °C in CA1 cells of GAD65-GFP slice cultures at DIV 13-19. Recordings were made on a Multiclamp 700B amplifier (Molecular Devices) and stored using pClamp 10 software. To isolate sIPSCs, 20 μM DNQX and 50 μM APV were added to the recording ACSF. For mIPSCs, 0.5 μM TTX was added as well. Thick walled borosilicate pipettes of 3-6 MΩ were filled with an internal solution containing 70 mM K-gluconate, 70 mM KCl, 0.5 mM EGTA, 10 mM HEPES, 4 mM MgATP, 0.4 mM NaGTP, and 4 mM Na_2_Phosphocreatine. Cells were excluded from analysis if the series resistance increased more than 35 %. IPSCs were automatically detected in Clampfit and further analyzed in custom Matlab scripts. Detected events within 3 ms of each other were merged and events smaller than 3 times the RMS of the signal were excluded. The cumulative distributions for individual experiments were interpolated to generate the average distribution.

### Immunohistochemistry, confocal imaging and image analysis

For *post hoc* immunohistochemistry, organotypic hippocampal slices were fixed in 4 % (w/v) paraformaldehyde for 30 minutes at room temperature. Slices were rinsed in phosphate buffer and permeabilized with 0.5 % TritonX-100 in phosphate buffer for 15 minutes. Slices were blocked with 0.2 % TritonX-100, 10 % goat serum (ab7481, Abcam) in phosphate buffer for 60 minutes. Primary antibodies were applied overnight at 4°C in blocking solution. After washing, slices were incubated with secondary antibodies in blocking solution for 4h at room temperature. Slices were washed and mounted on slides in Vectashield mounting medium (Vector Labs).

The following primary and secondary antibodies were used: rabbit α-VGAT (1:1000; Synaptic Systems, 131 003), mouse α-gephyrin (1:1000; Synaptic Systems, 147 011), guinea pig α-VGLUT (1:400; Millipore, AB5905), rabbit α-Homer (1:1000; Synaptic Systems, 160 002), mouse α-myc (1:100; Oncogene Research Products, OP10), mouse α-MET (1:500; Santa Cruz Biotechnology, sc-8057), Alexa405-, Alexa-488 and Alexa-568 conjugated secondary antibodies (Invitrogen). For staining MET we used a previously described myc-tagged nanobody, which was shown to recognize MET with low nanomolar affinity (Heukers et al., 2014). We visualized the nanobody with an antibody against the C-terminal myc tag. We validated the nanobody staining in primary hippocampal cultures using a previously described immunostaining protocol (Esteves da Silva et al., 2015).

For immunostainings, high resolution confocal laser scanning microscopy was performed on a Zeiss LSM-700 system with a Plan-Apochromat 63x 1.4 NA oil immersion objective. Each image was a z-series of 11-35 images (0.3 μm z step size), each averaged 4 times. The imaging area in the CA1 region was 78 x 78 μm (1024 x 1024 pixels). The confocal settings were kept the same to compare fluorescence intensities between slices.

For the quantification of VGAT and gephyrin intensities per image, we determined per image the mean intensity of 3 randomly chosen areas of 10 x 10 μm of the average projection image from the 5 middle z-layers. For the cumulative plots individual values (per area) were used. Synaptic puncta size and number were determined using the PunctaAnalyzer plugin, and inhibitory synapses were defined as overlapping VGAT and gephyrin puncta. For determining co-localization of GFP-labeled boutons with synaptic marker VGAT or with MET, we manually inspected individual boutons through all z-sections. A bouton was only considered positive when at least one z stack of the bouton overlapped with VGAT or MET staining. The images were median-filtered only for illustration purposes.

### Statistical Analysis

Data are represented as mean values ± standard error of the mean, unless stated otherwise. Statistical analysis was performed using GraphPad Prism software. Results from treatment and control experiments were compared using the Mann-Whitney U test (MW). The Chi-Square test (χ^2^) was used for comparing the fraction of axons with/without stabilizing boutons. For comparing multiple groups, we used the Kruskal-Wallis test (KW) followed by a posthoc Dunn’s comparison test. We used a One-Way ANOVA followed by a Dunnett’s multiple comparison test (One-Way ANOVA) to compare the effect of wash-in of PHA over time. We used a Two-Way ANOVA followed by a Sidak’s multiple comparisons test (TwoWay ANOVA) to compare treatment effects at multiple time points. For the comparison of cumulative distributions, we used the Kolmogorov-Smirnov (KS) test. We have indicated the tests and p-values in the figure legends. Differences between control and treatment were considered significant when *p* < 0.05 (*, *p* < 0.05; **, *p* < 0.01; ***, *p* < 0.001). In all figure legends and text, N indicates the number of independent experiments, and n indicates the number of axons/images analyzed.

## RESULTS

We performed time-lapse two-photon microscopy in organotypic hippocampal cultures from GAD65-GFP mice to monitor the dynamics of inhibitory boutons in the CA1 region of the hippocampus (Wierenga et al., 2008; Schuemann et al., 2013). In GAD65-GFP mice, approximately 20% of the CA1 interneurons express GFP. The majority of GFP-labeled interneurons express reelin and VIP, while parvalbumin and somatostatin expression is nearly absent (López-Bendito et al., 2004; Wierenga et al., 2010). High-resolution image stacks of GFP-labeled inhibitory axons were acquired every 10 minutes, for a total period of 150 minutes (15 time points). Inhibitory boutons were remarkably dynamic and many boutons appear, disappeared and reappeared during the course of the imaging period. To bias our analysis towards synaptic events, we only included boutons that appeared for at least 2 time points at the same location during the imaging period. We distinguished two main classes of boutons: persistent boutons, which were present during all time points, and non-persistent boutons, which were absent during one or more time points during the imaging session (Fig. 1A,B). Approximately 77 % (with standard deviation of 12 %) of inhibitory boutons at any given time point were persistent (Fig. 1C), and they reflect inhibitory synapses (Wierenga et al., 2008; Müllner et al., 2015). Non-persistent boutons reflect locations where inhibitory synapses are ‘in transition’, e.g. where synapses are being formed or disassembled (Wierenga et al., 2008; Dobie and Craig, 2011; Fu et al., 2012; Schuemann et al., 2013). Based on the presence or absence of non-persistent boutons during a baseline and wash-in period (details are given in the methods section), we distinguished 5 subgroups of non-persistent boutons: new (N; absent during baseline), lost (L; absent during wash-in), stabilizing (S; non-persistent during baseline, persistent during wash-in), destabilizing (D; persistent during baseline, non-persistent during wash-in) and transient (non-persistent in both periods). These different subgroups of non-persistent boutons not only differed in their incidence and duration (Fig. 1C,D), but also in their molecular composition, as assessed by immunostaining for the presynaptic vesicular GABA transporter (VGAT) and the postsynaptic scaffold gephyrin (Fig. 1E). Stabilizing boutons, which were present for at least 90 minutes before fixation, showed similar association with VGAT and gephyrin as persistent boutons, indicating that they are nascent inhibitory synapses that have started to recruit pre-and postsynaptic proteins within this period. Newly formed boutons, which were present for a short period before fixation, showed a lower percentage of VGAT and gephyrin association. Boutons with longer total lifetime before fixation showed higher association with VGAT and gephyrin, suggesting a gradual recruitment of proteins over the imaging period (Fig. 1F). Recruitment of gephyrin appeared delayed compared to VGAT, as previously reported (Wierenga et al., 2008; Dobie and Craig, 2011). These data demonstrate that inhibitory presynaptic boutons are dynamic structures that are continuously being formed and disassembled along the axons, and suggest that non-persistent boutons reflect boutons at different stages of inhibitory synapse assembly and disassembly.

### Inhibitory bouton stabilization during treatment with Sema4D

It was recently shown that class 4 semaphorin Sema4D can rapidly induce an increase of functional inhibitory synapses in hippocampal dissociated cultures (Kuzirian et al., 2013). However, these data could not resolve if Sema4D directly promotes synapse formation or rather prevents ongoing synapse elimination, thereby indirectly increasing synaptic density. To examine the effect of Sema4D on ongoing inhibitory bouton dynamics, we bath applied the extracellular domain of mouse Sema4D conjugated to the Fc region of mouse IgG2A (Sema4D; 1 nM) and compared inhibitory bouton dynamics during a baseline period of 5 time points and during Sema4D treatment in the subsequent 10 time points (Fig. 2A). We used Fc alone (1 nM) as a control treatment (Kuzirian et al., 2013). Bath application of Sema4D did not affect overall axonal morphology (Fig. 2A) and did not change the density of inhibitory boutons (Fig. 2B). However, when we analyzed the different subgroups of non-persistent boutons, we found that Sema4D treatment specifically enhanced the fraction of stabilizing boutons from 6 ± 2% to 16 ± 3 % (Fig. 2C). Indeed, this effect was also clear when we analyzed the absolute density of boutons. The density of stabilizing boutons was increased by >2-fold, while other subgroups of boutons were unaffected (Fig. 2D-H). To examine how Sema4D-induced stabilization developed over time, we quantified the number of boutons that were present for 5 consecutive time points during the baseline and the wash-in period. We found that Sema4D induced a marked increase in these boutons over the course of the wash-in period (% stabilization, Fig. 2I), and strongly enhanced the number of boutons that had stabilized at the end of this period (last 5 time points; Fig. 2J). Stabilizing boutons are relatively rare in our slices, as under control conditions only 40% of the axons display one or more stabilizing boutons. Treatment with Sema4D significantly increased this fraction to 77% (Fig. 2K). Altogether, these data show that Sema4D treatment in intact tissue specifically promotes the stabilization of inhibitory boutons within tens of minutes, without affecting synapse elimination.

**Figure 2.**
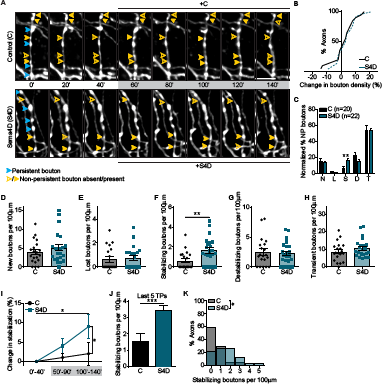
Sema4D treatment promotes inhibitory bouton stabilization. **(A)** Time-lapse two-photon images of GFP-labeled inhibitory axons in the CA1 region of the hippocampus during baseline (5 time points) and wash-in (10 time points; grey box) of 1 nM Fc - control (C; upper panel) or 1 nM Sema4D-Fc (S4D; bottom panel). Only every second image is shown for clarity. Persistent (blue) and non-persistent (yellow) boutons are indicated by arrowheads. Filled arrowheads indicate that the bouton is present, and empty arrowheads indicate that the bouton is absent at that time point. Images are maximum intensity projections of 11-18 z stacks. Time in minutes. Scale bar 5 μm.
**(B)** Cumulative distribution of the change in mean bouton density during the wash-in period compared to baseline after wash-in of C or S4D (MW, *p* = 0.83).
**(C)** Average fraction of subgroups of non-persistent boutons in C- and S4D-treated axons: N - new (MW, *p* = 0.86); L - lost (MW, *p* = 0.93); S - stabilizing (MW, *p* = 0.003); D - destabilizing (MW, *p* = 0.25); T - transient (MW, *p* = 0.89).
**(D-H)** Density of new (D; MW, *p* = 0.41), lost (E; MW, *p* = 0.61), stabilizing (F; MW, *p* = 0.003), destabilizing (G; MW, *p* = 0.84) and transient (H; MW, *p* = 0.34) boutons in axons treated with 1 nM Fc (C) and 1 nM Sema4D-Fc (S4D). Each dot represents an individual axon.
**(I)** Stabilization of inhibitory boutons, as determined by the change (compared to baseline) in density of boutons that were present at 5 consecutive time points during the imaging period: 0’-40’ (baseline), 50’-90’ (wash-in) and 100’-140’ (wash-in). Two-way ANOVA analysis showed a significant effect of both treatment (*p* = 0.04) and time (*p* = 0.03).
**(J)** Density of boutons that stabilized in the last 5 time points (TPs) (MW, *p =* 0.0008).
**(K)** Frequency distribution of the stabilizing bouton density in C- and S4D-treated axons (χ^2^, *p* = 0.03). Data are represented as mean ± SEM. Data from 20 control axons (N=6) and 22 S4D-treated axons (N=5).

### Sema4D-induced stabilization of inhibitory boutons is the first step of inhibitory synapse formation

We next assessed whether Sema4D-induced bouton stabilization also results in an increase of inhibitory synapses in our slices. We first examined if longer Sema4D treatment could enhance the bouton stabilization effect. We compared dynamics of individual boutons during baseline and after 6 h treatment (400 minutes total treatment) and found that longer Sema4D treatment also induced prominent bouton stabilization, measured as fraction as well as absolute density (Fig. 3A,B). However, the 6 h treatment did not increase bouton stabilization beyond the 2 h level (Fig. 3C), suggesting that only a limited number of inhibitory boutons can be stabilized by Sema4D treatment, resulting in saturation of the treatment effect already after 2 hr. In addition to promoting bouton stabilization, with longer treatments we also detected a small reduction in the fraction of transient boutons (Fig. 3A). This effect was only revealed by analyzing the changes in density over time (Fig. 3D), which suggest that this may reflect an indirect effect of prolonged bouton stabilization. These results indicate that the Sema4D-induced stabilization of inhibitory boutons persists, but does not further increase, with longer treatments.

**Figure 3.**
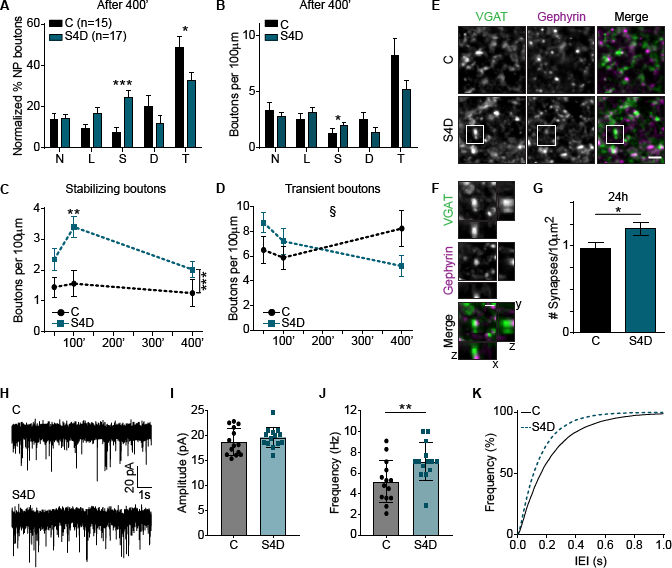
Sema4D increases overall inhibitory synaptic density. **(A)** Fraction of non-persistent boutons after treatment with 1 nM Fc (control; C) or 1 nM Sema4D-Fc (S4D) for 6 hours (400 minutes of total treatment). N - new (MW, *p* = 0.91) L - lost (MW, *p* = 0.13); S - stabilizing (MW, *p* = 0.0003); D - destabilizing (MW, *p* = 0.16); T - transient (MW, *p* = 0.02).
**(B)** Density of non-persistent boutons after treatment with 1 nM Fc (C) or 1 nM Sema4D- Fc (S4D) for 6 hours. N: MW, *p* = 0.74; L: MW, *p* = 0.29; S: MW, *p* = 0.03; D: MW, *p* = 0.09; T: MW, *p* = 0.11.
**(C)** Density of stabilizing boutons after treatment with Fc or S4D for 50, 100 and 400 minutes. Two-way ANOVA analysis showed that S4D increased bouton density independent of time (*p* = 0.0002). At 100’, *p* = 0.005 (Sidak’s multiple comparisons test).
**(D)** Same as B, but for transient boutons. Two-Way ANOVA analysis indicated a significant interaction between treatment and time (§; *p* = 0.03).
**(E)** Representative images of CA1 dendritic area of GAD65-GFP hippocampal slices treated with 1 nM Fc (C) or 1 nM Sema4D-Fc (S4D) for 24 h, and immunostained for VGAT (green) and gephyrin (magenta). Images are average intensity projections of 5 z stacks. Scale bar 2 μm.
**(F)** Example of an inhibitory synapse (white box in D), identified as the apposition of VGAT (green) and gephyrin (magenta) puncta. The respective xz and yz projections show the close apposition of the two markers. Images are maximum intensity projections of 6 z stacks. Scale bar 1 μm.
**(G)** Density of inhibitory synapses in slices treated with Fc or S4D for 24 h (MW, *p* = 0.03).
**(H)** Representative whole-cell voltage-clamp recordings of miniature inhibitory postsynaptic currents (mIPSCs) from CA1 pyramidal cells in organotypic hippocampal slices treated for 24 h with 1 nM Fc/DMSO (C) or 1 nM S4D/DMSO (S4D).
**(I-J)** Mean mIPSC amplitude (H) and frequency (I) in CA1 cells after 24 h treatment with Fc or S4D (H: MW*,p* = 0.35; I: MW*,p* = 0.008).
**(K)** Cumulative distribution of inter-event interval (IEI) of mIPSCs. Data are represented as mean ± SEM. Data in A,B from 15 control axons (N=4) and 17 S4D- treated axons (N=4), data in G from 15 control images (N=3) and 15 S4D images (N=3), and data in H-K from 14 control cells (N=5) and 14 S4D-treated cells (N=7).

We next asked if Sema4D-induced inhibitory bouton stabilization leads to the formation of functional synapses. We treated organotypic hippocampal slices with 1 nM Fc or 1 nM Sema4D for 24 h, and determined overall inhibitory synapse density by immunohistochemistry. We used antibodies against presynaptic VGAT and postsynaptic gephyrin to visualize inhibitory synapses (Fig. 3E,F). Sema4D induced a clear 24 ± 7 % increase in the density of inhibitory synapses (Fig. 3G), suggesting that the observed Sema4D-induced bouton stabilization after 2 h resulted in the formation of new synapses after 24 h. We used electrophysiological recordings to verify that these synapses were functional. In agreement with the immunohistochemistry results, we found that 24 h treatment with Sema4D increased the frequency of miniature inhibitory postsynaptic currents (mIPSCs) by 37 % (from 5.2 ± 0.5 to 7.1 ± 0.5 Hz), while mIPSC amplitude was not affected (Fig. 3H-K). To determine the time course of the recruitment of pre-and postsynaptic elements during synapse formation, we quantified VGAT and gephyrin immunostaining after 2, 6 and 24 h treatments. Treatment with Sema4D induced an increase in the area of VGAT puncta, without affecting their density (Fig. 4A-C). For gephyrin, Sema4D treatment caused an increase in puncta density, but not in their size (Fig. 4D-F). The average puncta intensity was not affected (at 24 h, VGAT: 107 ± 4 % of control, *p* = 0.35 (MW); gephyrin: 106 ± 5 % of control, *p* = 0.51 (MW)). Interestingly, the time course for presynaptic and postsynaptic changes was different. Whereas an increase in presynaptic VGAT area could already be detected after 6 h, the increase in postsynaptic gephyrin density was only evident after 24 h. These data are consistent with gradual increase in presynaptic vesicle content and subsequent acquisition of postsynaptic scaffolds at newly formed inhibitory synapses (Wierenga et al., 2008; Dobie and Craig, 2011). Together, these data indicate that the initial Sema4D-induced stabilization of inhibitory boutons is followed by a slower maturation process, resulting in an overall increase in functional inhibitory synapses after Sema4D treatment.

**Figure 4.**
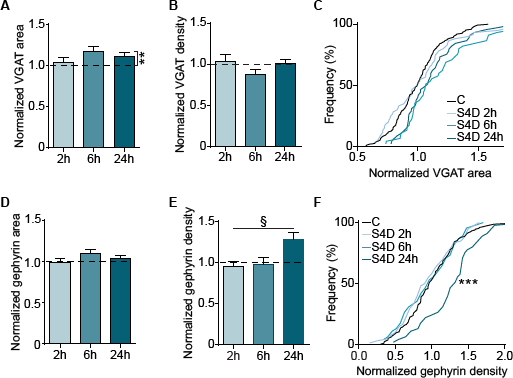
VGAT and gephyrin undergo different changes in response to Sema4D. **(A)** Normalized area of presynaptic vesicular GABA transporter (VGAT) puncta (after treatment with 1 nM S4D for 2 h, 6 h and 24 h). Dotted line represents control (treatment with 1nM Fc for 2 h, 6 h and 24 h). Two-way ANOVA analysis showed that S4D treatment increased VGAT area independent of time (*p =* 0.005).
**(B)** Normalized density of VGAT puncta, after treatment with 1 nM S4D for 2 h, 6 h and 24 h. Dotted line represents control (treatment with 1nM Fc for 2 h, 6 h and 24 h).
**(C)** Cumulative distributions of the normalized area of VGAT after treatment with 1 nM S4D for 2, 6 and 24 h. Black line represents the normalized control values. *p* = 0.81, *p* = 0.08 and *p* = 0.14 (KS) for 2, 6 and 24 h, respectively.
**(D-E)** Same as in A-B, but for normalized area (D) and density (E) of postsynaptic gephyrin puncta. Two-way ANOVA analysis showed a significant effect of time (*p* = 0.04) and an interaction between treatment and time (§; *p* = 0.04) in E.
**(F)** Same as in C, but for normalized gephyrin density. *p* = 0.99, *p* = 0.99 and *p* < 0.0001 (KS) for 2, 6 and 24 h, respectively. Data are represented as mean ± SEM. Data from 15-20 control images (N=3-4) and 15-20 S4D images (N=3-4) per time point.

### Sema4D-induced bouton stabilization relies on network activity

We previously showed that inhibitory bouton dynamics are regulated by neuronal activity (Schuemann et al., 2013). We therefore asked whether Sema4D-induced stabilization of inhibitory boutons depended on network activity. Blocking activity by bath application of tetrodotoxin (TTX) slightly decreased overall bouton dynamics in our slices (data not shown), which is in accordance with our previous findings (Schuemann et al., 2013). However, we found that in the presence of TTX Sema4D treatment no longer induced stabilization of inhibitory boutons, and that Sema4D treatment even led to a reduction in bouton stabilization compared to control (Fig. 5A,B). Indeed, whereas under control conditions Sema4D treatment increased the number of axons that displayed stabilizing boutons, it led to a decrease in the presence of TTX (Fig. 5C,D). These findings demonstrate that Sema4D treatment affects bouton dynamics in an activity-dependent manner, and indicate that Sema4D promotes the stabilization of inhibitory presynaptic boutons only in active neuronal networks.

**Figure 5.**
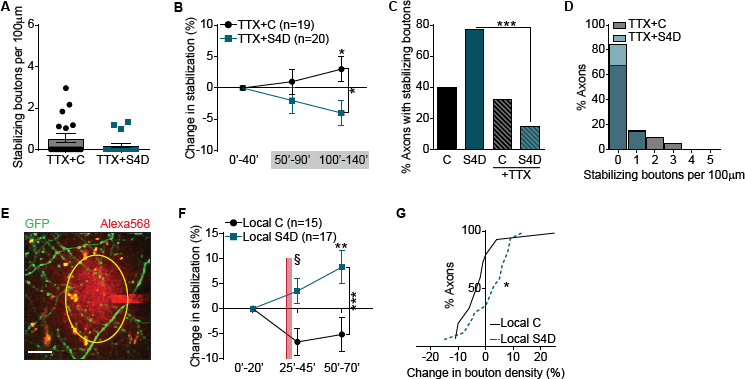
Sema4D induces local stabilization of inhibitory boutons. **(A)** Density of stabilizing boutons in axons treated with 1 nM Fc (C) and 1 nM Sema4D-Fc (S4D), in the presence of 0.5 μM TTX (MW, *p* = 0.17).
**(B)** Stabilization of inhibitory boutons upon treatment with C or S4D in the presence of 0.5 μM TTX, determined by the change (compared to baseline) in density of boutons that were present at 5 consecutive time points during the imaging period: 0’-40’ (baseline), 50’-90’ (wash-in) and 100’-140’ (wash-in). Two-way ANOVA analysis showed a significant effect of treatment (*p* = 0.01). At 100’-140’, *p* = 0.04 (Sidak’s multiple comparisons test).
**(C)** Fraction of axons with stabilizing boutons in axons treated with C or S4D, in normal or activity-depleted slices with TTX (χ^2^(p-values are Bonferroni-corrected): C vs S4D, *p* = 0.01; C vs C+TTX, *p* = 0.58; C+TTX vs S4D+TTX, *p* = 0.22; S4D vs S4D+TTX, *p* < 0.0001).
**(D)** Frequency distribution of the stabilizing bouton density in C- and S4D-treated axons, in the presence of 0.5 μM TTX (x, *p* = 0.17).
**(E)** Representative image of the local treatment of GFP-labeled inhibitory axons in the CA1 region of the hippocampus. The pipette was filled with Alexa568 (red) to visualize the area of the puff (yellow circle). Scale bar 10 μm.
**(F)** Same as B, but for local treatment with 10 nM Fc (control, C) or 10 nM S4D. Red line marks the puffing. Two-way ANOVA analysis showed a significant effect of treatment (*p* = 0.0002) and an interaction between treatment and time (§; *p* =0.02). At 50’-70’, *p* = 0.003 (Sidak’s multiple comparisons test).
**(G)** Cumulative distribution of the change in mean bouton density after local treatment with C or S4D compared to baseline (MW, *p* = 0.045). Data are represented as mean ± SEM. Data in A-D from 19 control axons (N=5) and 20 S4D-treated axons (N=5), and in F-G from 15 control axons (N=6) and 17 S4D-treated axons (N=6).

### Local Sema4D-induced bouton stabilization

Under physiological circumstances, Sema4D is a membrane-attached protein acting locally (Pasterkamp, 2012; Raissi et al., 2013). Presynaptic boutons along the same axon interact and share presynaptic proteins and vesicles (Staras, 2007; Bury and Sabo, 2016) and we wondered if local Sema4D signaling would act differently compared to ubiquitous activation of Sema4D signaling during bath application. We therefore locally applied Sema4D to short stretches (~40 p,m) of inhibitory axons (Fig. 5E). Local application with control solution appeared to slightly reduce local bouton stabilization (compare control curves in 5F and 2I), possibly from mechanical pressure. In contrast, local application of Sema4D induced robust stabilization of inhibitory boutons in these axons (Fig. 5F), resulting in a significant increase in local bouton density in these short axon stretches (Fig. 5G). This indicates that local application of Sema4D is more potent to induce axonal changes than bath application, which failed to induce a change in overall bouton density (compare Fig. 2B). This suggests that stabilizing boutons may compete for presynaptic components within individual axons when Sema4D is bath applied, limiting overall bouton density. Together, our results demonstrate that Sema4D-signaling is capable to mediate rapid changes in local bouton density of inhibitory axons in an activity-dependent manner.

### Actin remodeling by low doses of LatrunculinB promotes stabilization of inhibitory boutons

The Sema4D effect on inhibitory synapses was previously shown to be dependent on its receptor PlexinB1 (Kuzirian et al., 2013). Sema4D/PlexinB1 signaling induces changes in the intracellular actin cytoskeleton via multiple small GTPase signaling pathways in many different cell types (Zhou et al., 2008; Cagnoni and Tamagnone, 2014). Some of these downstream signaling pathways, which can modify actin in multiple ways, are mediated by receptor tyrosine kinases, such as MET and ErbB-2, acting as co-receptors for PlexinB1. It was shown in breast carcinoma cells that, when MET is co-activated, Sema4D/PlexinB1 signaling reduces RhoA levels and this results in actin depolymerization, while co-activation of ErbB-2 leads, via RhoA activation, to actin polymerization (Swiercz et al., 2008; Sun et al., 2012). To examine how the actin cytoskeleton is involved in inhibitory bouton dynamics, we studied the effect of two actin remodeling drugs in our system with intended opposite effects: the actin monomer sequestering drug LatrunculinB (LatB), which is generally considered an actin depolymerizing drug, and the actin filament stabilizer Jasplakinolide (Jasp), which promotes actin polymerization. In the low concentrations that we use here (100 nM LatB and 200 nM Jasp) these drugs perturb the actin cytoskeleton without affecting synaptic function (Honkura et al., 2008; Rex et al., 2009). None of the treatments changed overall axon morphology (Fig. 6A). We found that the fraction of stabilizing boutons was increased in the presence of LatB, but not in the presence of Jasp (Fig. 6B,C). The effect of LatB seemed highly specific for stabilizing boutons, as the other bouton subgroups were not affected. Indeed, we found that LatB specifically increased the absolute density of stabilizing boutons by almost 2-fold (Fig. 6D) and increased the fraction of axons with stabilizing boutons (Fig. 6E). The rapid and highly specific action of LatB suggests a direct action on the local actin cytoskeleton. The changes in bouton dynamics after LatB treatment were surprisingly similar to Sema4D treatment (Fig. 2F and 2K). Our findings suggest that inhibitory bouton dynamics are regulated by specific changes in the actin cytoskeleton and that conditions favoring actin depolymerization promote bouton stabilization.

**Figure 6.**
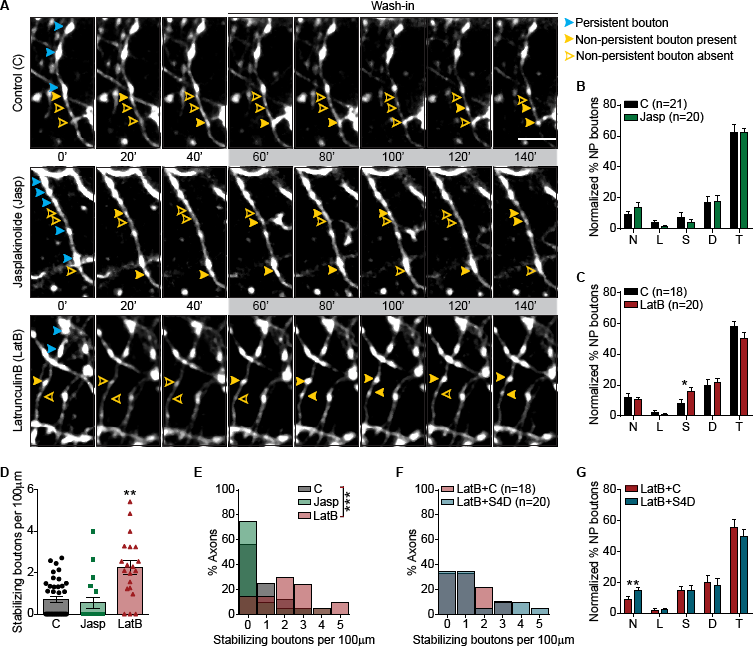
Inhibitory bouton dynamics are regulated by actin. **(A)** Time-lapse two-photon images of GAD65-GFP-labeled axons in the CA1 region of the hippocampus during baseline (5 time points) and wash-in (10 time points; grey box) of DMSO - control (C; upper panel), 200 nM Jasplakinolide (Jasp; middle panel) or 100 nM LatrunculinB (LatB; bottom panel). Only every second image is shown for clarity. Persistent and non-persistent boutons are indicated as in Figure 2. Images are maximum intensity projections of 12-14 z stacks. Time in minutes. Scale bar 5 μm.
**(B)** Fraction of non-persistent (NP) boutons in C and Jasp-treated axons: N - new (MW, *p* = 0.37); L - lost (MW, *p* = 0.18); S - stabilizing (MW, *p* = 0.49); D - destabilizing (MW, *p* = 0.95); T - transient (MW, *p* = 0.93).
**(C)** Same as in B, but for C and LatB-treated axons (N: MW, *p* = 0.99; L: MW, *p* = 0.66; S: MW, *p* = 0.01; D: MW, *p* = 0.6; T: MW, *p* = 0.29).
**(D)** Density of stabilizing boutons in control, Jasp- (MW: *p* = 0.55) and LatB-treated axons (MW: *p* = 0.005).
**(E)** Frequency distribution of the stabilizing bouton density in C, Jasp- and LatB-treated slices (χ^2^; C vs Jasp, *p* = 0.31; C vs LatB, *p* = 0.0005).
**(F)** Same as E, but for combined treatment with 100 nM LatB/1 nM Fc (LatB+C) or 100 nM LatB/1 nM Sema4D (LatB+S4D) (χ^2^,*p* = 0.37).
**(G)** Same as B, but for combined treatment with LatB+C or LatB+S4D (N: MW, *p* = 0.005; L: MW, *p* = 0.58; S: MW, *p* = 0.96; D: MW, *p* = 0.82; T: MW, *p* = 0.52). Data are represented as mean ± SEM. Data in B from 21 control axons (N=6) and 20 Jasp-treated axons (N=5), in C from 18 control axons (N=5) and 20 LatB-treated axons (N=5) and in F-G from 18 LatB+Fc- (N=4) and 20 LatB+S4D-treated axons (N=5).

The similarity between stabilization of inhibitory boutons induced by treatment with LatB or Sema4D suggests that both treatments may induce a similar effect on intracellular actin. As we found that only a specific subset of inhibitory boutons were stabilized by Sema4D treatment (Fig. 2C), we wondered if these were the same boutons that responded to LatB. To test this, we treated slices with a combination of LatB and Fc or LatB and Sema4D. We found that bouton stabilization by LatB occluded a further increase by co-application with Sema4D (Fig. 6F and 6G), although it did increase the fraction of new boutons. These results suggest that LatB and Sema4D treatment act to stabilize a specific, overlapping, subset of inhibitory boutons.

We then wondered if treatment with the actin depolymerizing drug LatB would be sufficient to induce inhibitory synapse formation, similarly to the Sema4D treatment. Interestingly, we observed that although LatB induced changes in VGAT and gephyrin puncta after 2 h (Fig. 7A-F), these changes were not coordinated and did not result in an increase in the density of inhibitory synapses (Fig. 7G). Gephyrin and VGAT staining returned to baseline with longer LatB treatment. Together, our data suggest that LatB and Sema4D induce rapid stabilization of the same subgroup of inhibitory boutons, but that only Sema4D signaling leads to coordinated pre- and postsynaptic changes resulting in inhibitory synapse formation. This indicates that presynaptic bouton stabilization alone is not enough to induce inhibitory synapse formation and that additional signaling may be required.

**Figure 7.**
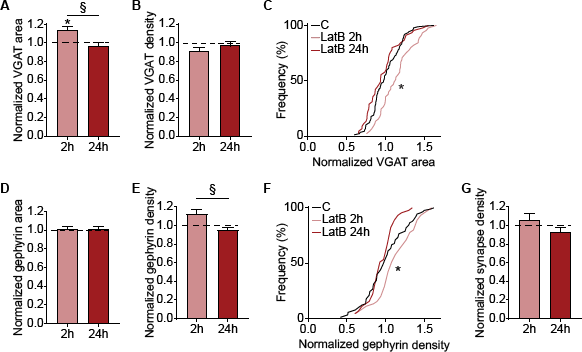
Latrunculin B treatment does not promote inhibitory synapse formation. **(A)** Normalized area of presynaptic vesicular GABA transporter (VGAT) puncta (after treatment with 100 nM LatB for 2 h and 24 h). Dotted line represents control (treatment with DMSO for 2 h and 24 h). Two-way ANOVA analysis indicated a significant effect of time (*p*= 0.02) and an interaction between treatment and time (§, *p* = 0.02).
**(B)** Normalized density of VGAT, after treatment with 100 nM LatB for 2 h and 24 h. Dotted line represents control (treatment with DMSO for 2 h and 24 h).
**(C)** Cumulative distributions of the normalized area of VGAT after treatment with 100 nM LatB for 2 and 24 h. Black line represents the normalized control values. *p* = 0.047 and 0.33 (KS) for 2 and 24 h, respectively.
**(D-E)** Same as in A-B, but for the area (D) and density (E) of postsynaptic gephyrin puncta. In E, Two-way ANOVA analysis showed a significant effect of time (*p* =0.04) and interaction between treatment and time (§, *p* =0.04).
**(F)** Same as in I, but for normalized gephyrin density. *p* = 0.047 and 0.33 (KS) for 2 and 24 h, respectively.
**(H)** Same as A, but for normalized density of inhibitory synapses. Data are represented as mean ± SEM. Data from 15 control images (N=3) and 15 LatB images (N=3) per time point.

### Inhibitory bouton stabilization by Sema4D requires MET

Our observation that LatB could mimic the Sema4D-induced stabilization of inhibitory boutons points to a possible involvement of MET in this process. We therefore assessed if MET activation is necessary for the observed Sema4D-induced stabilization of boutonsby making use of the highly specific MET inhibitor PHA-665752 (PHA) (Christensen et al., 2003; Deguchi et al., 2016). We first verified that adding PHA alone did not affect bouton dynamics (Fig. 8B) or spontaneous inhibitory postsynaptic currents (data not shown), indicating that MET is not very active under baseline conditions in our slices. Next, we treated our slices with Sema4D to induce bouton stabilization and compared bouton dynamics in the presence or absence of PHA (Fig. 8A, C-D) Blocking MET with PHA completely abolished the Sema4D-induced increase in the density of stabilizing boutons (Fig. 8D). In fact, blocking MET in combination with Sema4D treatment almost entirely abolished the occurrence of stabilizing boutons on our slices (Fig. 8E), while the other bouton subgroups were not much affected (8C). Consistent with the live imaging data, inhibiting MET with PHA also blocked the increase in VGAT staining intensity (Fig. 8F,G) and mIPSC frequency (Fig. 8H) after Sema4D treatment. Taken together, these data indicate that activation of MET is required for the Sema4D-induced stabilization of inhibitory boutons.

**Figure 8.**
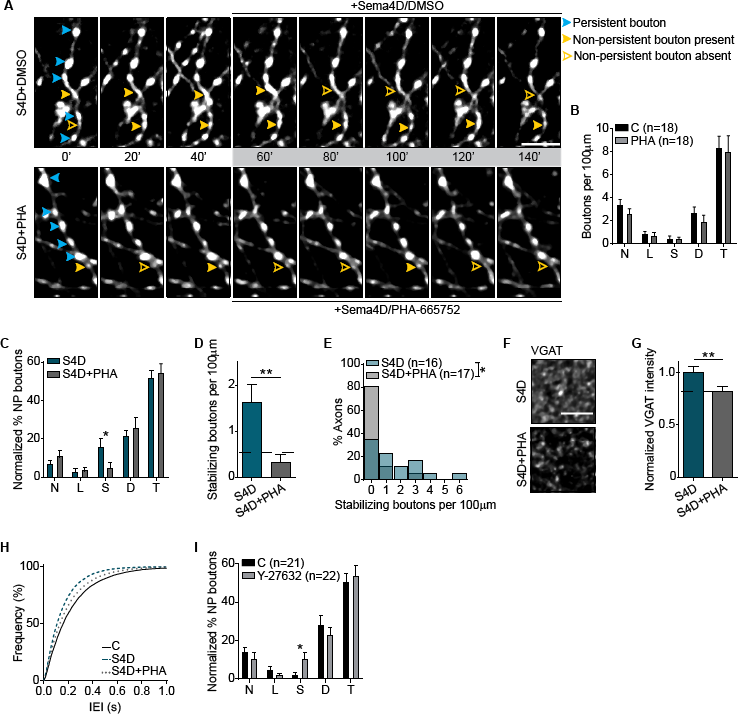
Inhibitory bouton stabilization by Sema4D requires MET. **(A)** Time-lapse two-photon images of GAD65-GFP-labeled axons in organotypic hippocampal slices during wash-in (grey box) of combination of 1 nM Sema4D and DMSO (S4D; upper panel) or combination of 1 nM Sema4D with 1 μM PHA-665752 (S4D+PHA; bottom panel). Only every second image is shown for clarity. Persistent and non-persistent boutons are indicated as in Figure 2. Images are maximum intensity projections of 15-16 z stacks. Scale bar 5 μm.
**(B)** Density of non-persistent boutons in slices treated with DMSO (C) and 1 μM PHA-665752 (PHA): N - new (MW, p = 0.28); L - lost (MW, p = 0.77); S - stabilizing (MW, p = 0.98); D - destabilizing (MW, p = 0.24); T - transient (MW, p = 0.67).
**(C)** Fraction of non-persistent (NP) boutons in S4D- and S4D+PHA-treated axons: N: MW, p = 0.34; L: MW, p = 0.74; S: MW, p = 0.01; D: MW, p = 0.64; T: MW, p = 0.53.
**(D)** Density of stabilizing boutons in slices treated with S4D or S4D+PHA (MW, p = 0.006). Dotted line represents control values.
**(E)** Frequency distribution of the stabilizing bouton density in S4D- and S4D+PHA- treated slices (χ^2^, *p* = 0.048).
**(F)** Representative images of hippocampal slices treated with S4D (upper panel) or S4D+PHA (bottom panel) for 100’, and stained for presynaptic VGAT. Images are average intensity projections of 5 z stacks. Scale bar 5 μm.
**(G)** Normalized mean staining intensity for VGAT in S4D- and S4D+PHA-treated slices (MW, p = 0.009). Control value is indicated with dotted line.
**(H)** Cumulative distribution of inter-event interval (IEI) of mIPSCs from CA1 pyramidal cells in organotypic slices after treatment with 1 nM Fc/DMSO (C), 1 nM S4D/DMSO (S4D) or 1nM S4D/1 μM PHA-665752 (S4D+PHA) for 24 h. C and S4D as in Figure 3K.
**(I)** Fraction of non-persistent boutons in axons treated with MQ (control) or 10 μM Y- 27632 (ROCK inhibitor): N: MW, *p* = 0.05; L: MW, *p* = 0.39; S: MW, *p* = 0.02; D: MW, *p* = 0.38; T: MW, *p* = 0.78. Data are represented as mean ± SEM. Data in B from 18 control axons (N=4) and 18 PHA-treated axons (N=4), in C-E from 17 S4D-treated axons (N=4) and 16 S4D+PHA-treated axons (N=4), in F-G from 16 images of S4D-treated slices (N=3) and 23 images of S4D+PHA-treated slices (N=4); in H from 14 control cells (N=5), 14 S4D-treated cells (N=7) and 17 S4D+PHA-treated cells (N=5), and in I from 21 control axons (N=5) and 22 Y-27632-treated axons (N=5).

As the actin depolymerization pathway downstream of Sema4D/PlexinB1 signaling via MET was previously shown to reduce intracellular RhoA activity (Swiercz et al., 2008; Sun et al., 2012), we tested if stabilization of inhibitory boutons could also be achieved by directly reducing ROCK activity, a well-known downstream effector of RhoA (Amano et al., 2010). We found that reducing ROCK signaling with the specific ROCK inhibitor Y-27632 also resulted in an increase in the density of stabilizing boutons in our slices (Fig. 8I), which was similar to the effect of LatB and Sema4D treatments. This suggests that the intracellular pathway that is activated by Sema4D/PlexinB1 signaling to induce stabilization of inhibitory boutons involves activation of MET receptor tyrosine kinase and reduction of ROCK activity to promote specific changes in the actin cytoskeleton.

### MET is enriched at a subset of inhibitory presynaptic boutons

Our pharmacological experiments do not address if Sema4D-induced changes in actin occur at the pre- or postsynaptic compartment. The subcellular localization of Sema4D and PlexinB1 is not known (Paradis et al., 2007), but the localization of MET has been described. Interestingly, it was reported that in postnatal tissue the majority of MET is localized in axons (Judson et al., 2009) and detailed EM analysis showed clusters of MET in the shaft of unmyelinated axons and in small presynaptic terminals (Eagleson et al., 2013). The majority of these terminals are glutamatergic (Tyndall and Walikonis, 2006; Xie et al., 2016), but possible MET expression in GABAergic axons was never addressed directly. To address the localization of MET in our slices, we made use of an antibody (Qiu et al., 2014) and a nanobody (Heukers et al., 2014) with demonstrated specificity for MET. We first confirmed that both label synapses in primary hippocampal cultures (Fig. 9A-C). The majority of MET puncta overlapped with excitatory synapses (Fig. 9A,B) in line with previous reports (Tyndall and Walikonis, 2006; Eagleson et al., 2013; Xie et al., 2016). However, clear association of MET with inhibitory presynapses was also observed in these cultures (Fig. 9B,C). We then used the MET nanobody and antibody to label MET in our hippocampal slices of GAD65-GFP mice (Fig. 9D). Although there was a quantitative difference, presumably reflecting a difference in labeling affinity, both methods clearly showed that a subset of GFP-labeled inhibitory boutons was enriched for MET (Fig. 9E). Comparison between the MET staining pattern with staining for postsynaptic gephyrin (compare Figs. 9F and 3F) suggests a presynaptic localization of MET at these inhibitory synapses, as MET puncta were often completely embedded in the GFP-labeled bouton. These data suggest that MET may be present in inhibitory axons and terminals to mediate Sema4D/plexinB1 signaling.

**Figure 9.**
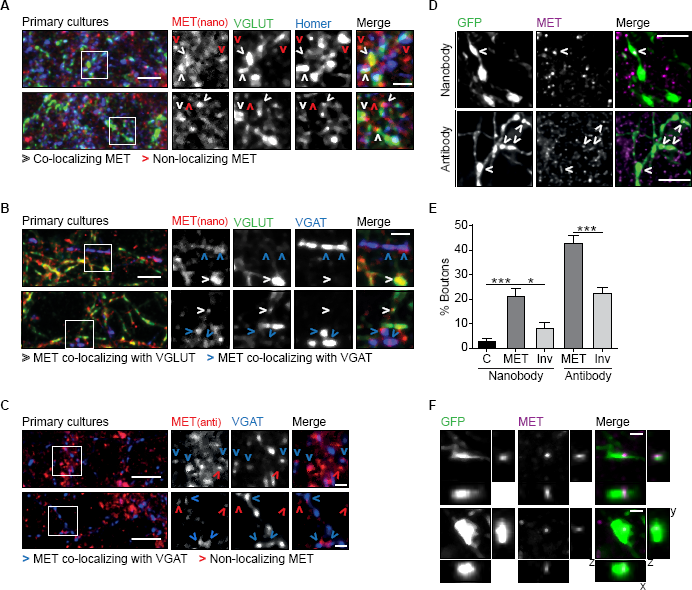
MET is enriched in a subset of inhibitory presynaptic boutons. **(A)** Images of primary cultures of hippocampal neurons immunostained with MET nanobody (red) and markers for excitatory synapses: presynaptic vesicular glutamate transporter (VGLUT; green) and postsynaptic Homer (blue). The majority of MET puncta co-localize with one or both markers (white arrows), but some MET puncta do not co-localize (red arrows). Images are maximum intensity projections of 13 stacks. Scale bar 5 μm (overview) and 2 μm (zoom).
**(B)** Same as A, but neurons were stained with MET nanobody (red) and markers for excitatory presynapses (VGLUT; green) and inhibitory presynapses (vesicular GAB A transporter VGAT; blue). White arrows indicate MET co-localizing with VGLUT and blue arrows indicate MET co-localizing with VGAT. Images are maximum intensity projections of 12 stacks. Scale bar 5 μm (overview) and 2 μm (zoom).
**(C)** Same as A, but hippocampal neurons were stained with MET antibody (red) and VGAT (blue). Blue arrows highlight MET puncta co-localizing with VGAT, while red arrows indicate MET puncta that do not co-localize with VGAT. Images are maximum intensity projections of 17-21 stacks. Scale bar 10 μm (overview) and 2 μm (zoom).
**(D)** Representative images of GFP-labeled inhibitory boutons (green) in hippocampal slices, stained with a nanobody (upper) and an antibody (lower panels) against MET (magenta). Images are maximum intensity projections of 5-6 z stacks. White arrows indicate MET enrichment in GFP-labeled boutons. Scale bar 5 μm.
**(E)** Fraction of GFP boutons positive for MET. Aspecific staining was determined by anti-myc staining without nanobody (‘C’; black) and random co-localization was determined by inverting the MET channel (‘Inv’; light gray) (Nanobody: KW, *p* = 0.002; Antibody: MW, *p* < 0.0001).
**(F)** Example of two inhibitory boutons (green) in hippocampal slices showing enrichment in MET (magenta), and the respective xz and yz projections. Images are maximum intensity projections of 6 z stacks. Scale bar 1 μm. Data are represented as mean ± SEM. Data in F from 10 control images (N=2), 12 images in MET and inverted group (N=3) for the nanobody staining and 15 images in MET and inverted group (N=3) for the antibody staining.

## DISCUSSION

By monitoring individual boutons over time in live brain slices, we observed that the primary action of Sema4D signaling is to stabilize presynaptic boutons of inhibitory axons within tens of minutes. These stabilizing boutons develop into mature, functional inhibitory synapses over the course of several hours. They rapidly acquire presynaptic vesicles as evidenced by an increase in VGAT staining, while recruitment of postsynaptic gephyrin was slower. We demonstrate for the first time that Sema4D-induced bouton stabilization is activity-dependent and that Sema4D signaling can induce local changes in bouton density. We found that inhibitory axons respond differently to Sema4D signaling in active and inactive networks (Fig. 5). These results suggest that inhibitory axons can respond very rapidly to local signals in their environment, but that the response is modulated by the state of the network and/or the local internal state of the axon. It was previously shown that Sema4D in inhibitory synapses signals via the PlexinB1 receptor. We now show that this signaling pathway requires coactivation of the receptor tyrosine kinase MET. The downstream intracellular pathway involves specific remodeling of the actin cytoskeleton, which can be mimicked by bath application of low levels of the actin depolymerizing drug LatB or by reducing ROCK activity. Our immunohistochemistry data suggest that MET is localized to the presynaptic compartment of GABAergic synapses. Our data is the first to show a role for the autism-linked MET in inhibitory synapses.

Our live imaging experiments give unique insight in the dynamics of inhibitory synapse formation in brain slices, which remain undetected with methods using stationary comparisons before and after treatment. In our slices, the majority of GFP-labeled boutons are persistent and display pre- and postsynaptic markers of mature inhibitory synapses, but a significant portion (~20-25%) of inhibitory boutons are non-persistent and represent locations where inhibitory synapses are ‘in transition’. At these axonal locations, inhibitory synapses are formed and disassembled in an apparent trial-and-error fashion (Wierenga et al., 2008; Dobie and Craig, 2011; Fu et al., 2012; Schuemann et al., 2013; Wierenga, 2017). We found that Sema4D signaling did not induce formation of inhibitory synapses *de novo,* but specifically stabilized boutons at locations where a bouton had occurred before. We also observed that Sema4D-induced bouton stabilization was not further enhanced by longer treatment (>2 h) or by co-application with LatB, suggesting that the number of boutons susceptible to Sema4D at any given time is limited. This suggests that Sema4D signaling is involved only at a specific stage during synapse formation and that boutons which are more mature or too immature do not respond to Sema4D. In primary cultures, a larger fraction of synapses are immature compared to intact tissue (Dobie and Craig, 2011; Kuriu et al., 2012), which may explain why the Sema4D effect on inhibitory synapses is stronger in primary neurons (Kuzirian et al., 2013). In our slices, Sema4D treatment increased inhibitory synapse density by ~20% after 24 hours (Fig. 3G), which is comparable to experience-dependent changes in inhibitory synapses observed *in vivo* (Keck et al., 2011; Chen et al., 2015; Villa et al., 2016).

One of the key observations of this study is that the primary action of Sema4D is to stabilize presynaptic boutons of inhibitory axons within tens of minutes. These stabilizing boutons develop into mature, functional inhibitory synapses over the course of several hours. It was previously shown that inhibitory synapses can be induced by postsynaptic gephyrin clustering (Flores et al., 2015), and rapid formation of new gephyrin clusters was observed after Sema4D treatment in primary cultures (Kuzirian et al., 2013), suggesting that Sema4D may promote inhibitory synapse formation via a postsynaptic mechanism. However, our data clearly show that gephyrin clustering after Sema4D treatment is delayed and that the primary action of Sema4D signaling is presynaptic bouton stabilization, arguing against a triggering mechanism via postsynaptic gephyrin. The increase in VGAT signal reflects recruitment of synaptic vesicles to newly forming synapses (Dobie and Craig, 2011; Schuemann et al., 2013). Bouton stabilization and gephyrin clustering were also induced by LatB, but LatB failed to induce new inhibitory synapses. This suggests that Sema4D signaling coordinates pre- and postsynaptic changes at emerging inhibitory synapses. The faster time course for the increase in postsynaptic gephyrin clusters in primary cultures (Kuzirian et al., 2013) may reflect an overall difference in neuronal maturation level. In young neurons, new gephyrin clusters can be rapidly induced by local GABA signaling (Oh et al., 2016), while in mature neurons prolonged or additional signaling may be required.

It was previously shown that Sema4D acts as a postsynaptic protein and requires PlexinB1 for promoting inhibitory synapse formation (Kuzirian et al., 2013; Raissi et al., 2013). Our data shows that this signaling pathway requires co-activation of MET, suggesting that the receptor tyrosine kinase MET acts as a co-receptor of PlexinB1 (Swiercz et al., 2008). PlexinB1 and MET receptors can form a complex which, upon Sema4D stimulation, results in cross-activation of both receptors (Giordano et al., 2002). It is currently not known if the PlexinB1 receptors that mediate the Sema4D signaling are located in the pre-or postsynaptic membrane and our data does not address this issue directly. However, our immunohistochemistry data (Fig. 9) suggest that MET receptors are localized in a subset of inhibitory synapses, in primary hippocampal cultures and organotypic slices. A presynaptic location of MET in inhibitory boutons suggests retrograde signaling of postsynaptic Sema4D via presynaptic plexinB1 receptors. Retrograde semaphorin signaling was recently demonstrated in Drosophila neuromuscular junction (Orr et al., 2017). However, cell-specific genetic studies will be needed to rule out a contribution of Sema4D signaling via postsynaptic receptors in inhibitory synapse formation.

Our data indicates that inhibitory bouton stabilization by activation of the Sema4D/PlexinB1 signaling pathway is induced through actin remodeling (Swiercz et al., 2008; Sun et al., 2012). The induced changes in actin are highly specific and not due to a general decrease in actin dynamics since Jasplakinolide did not affect inhibitory boutons. Treatment with the actin depolymerizing drug LatB or the ROCK inhibitor Y-27632 promoted bouton stabilization in a similar way as Sema4D, presumably by inducing similar changes in presynaptic actin at stabilizing boutons. Given that actin is abundantly present in all cells, it is surprising that bath application of LatB specifically promotes stabilization of immature boutons without affecting other inhibitory boutons. It is important to note that low doses of monomer sequestering drugs, such as LatB, do not lead to the complete disassembly of actin structures and leave postsynaptic spines and synaptic transmission intact (Honkura et al., 2008; Rex et al., 2009; Bleckert et al., 2012). Instead, LatB treatment may result in limited availability of actin monomers in small cellular compartments, which indirectly affects actin-regulating factors resulting in structural changes of the actin cytoskeleton (Ganguly et al., 2015; Suarez et al.,2015). Our data therefore suggest that the actin cytoskeleton at stabilizing boutons is different from other compartments and specifically sensitive to LatB. However, we stress here that our experiments cannot distinguish between pre- or postsynaptic effects of LatB, but the rapid (boutons are stabilized within 10 minutes) and highly specific effect seems to suggest a local action of LatB. Inhibitory synapses are usually localized directly on the dendritic shaft and not much is known about a possible role of postsynaptic actin structures at these synapses. Within axons, several actin-based structures have been described (Leterrier et al., 2017) and presynaptic actin has been implicated in transmission, plasticity, as well as synapse formation (Cingolani and Goda, 2008; Chia et al., 2013). In *C. elegans* and *Drosophila* it has been demonstrated that presynaptic actin structures undergo important remodeling during synapse formation (Chia et al., 2014; Piccioli and Littleton, 2014). It is currently not known which actin-regulating factors are involved in presynaptic bouton stabilization, but promising candidates include cortactin (Alicea et al., 2017), cofilin (Piccioli and Littleton, 2014) and Mical (Orr et al., 2017). Future studies will be necessary to unravel precise actin structures in mature and immature boutons and the role of actin-regulating factors during synapse formation.

Changes in inhibitory synapses play an important role in the rewiring of circuits during development and in response to behavioral demands during adulthood (Keck et al., 2011; Chen et al., 2015; Froemke, 2015) and defects in GABAergic synapses are associated with neurodevelopmental diseases (Hensch, 2004; Marín, 2012). Mutations in the MET gene are an established risk factor for autism spectrum disorder (ASD), as determined by various human imaging and genetic studies (Peng et al., 2013). It is a multifunctional receptor involved in many cellular pathways, and its exact role in ASD is not yet understood (Eagleson et al., 2017). Previous studies in neurons have implicated MET in regulating postsynaptic strength in excitatory neurons (Qiu et al., 2014; Lo et al., 2016), excitatory synapse formation (Xie et al., 2016) and interneuron migration (Martins et al., 2011). Our data demonstrate that activation of MET is also an essential part of the Sema4D-signaling pathway promoting activity-dependent inhibitory bouton stabilization, indicating a novel role of MET in the assembly of inhibitory presynapses.

## Acknowledgements

We would like to thank G. Szábo for kindly providing the GAD65-GFP mice, R. van Dorland for technical support, M. van Kesteren for analysis of LatB immunodata, S. Paradis for helpful comments and scientific discussions and A. Akhmanova, G.G. Turrigiano and R.J. Pasterkamp for critically reading the manuscript. This work was supported by the People Programme (Marie Curie Actions) of the European Union’s Seventh Framework Programme FP7/2007-2013/ under REA grant agreement 289581 (C.P.F.), a Marie Curie Reintegration Grant 256284 (C.J.W.) and the Netherlands Organization for Scientific Research (NWO-VIDI, C.J.W., NWO-VICI, C.C.H.).

## Notes

**Conflict of Interest** The authors declare no competing financial interests.

